# Lateral organ diversification in plants mediated by the ALOG protein family of transcription factors

**DOI:** 10.1101/689109

**Authors:** Satoshi Naramoto, Victor Arnold Shivas Jones, Nicola Trozzi, Mayuko Sato, Kiminori Toyooka, Masaki Shimamura, Sakiko Ishida, Kazuhiko Nishitani, Kimitsune Ishizaki, Ryuichi Nishihama, Takayuki Kohchi, Liam Dolan, Junko Kyozuka

## Abstract

Land plant shoot structures evolved a diversity of lateral organs as morphological adaptations to the terrestrial environment, in which lateral organs independently evolved in each lineage in the sporophyte or gametophyte generation. The gametophyte meristem of the basally-diverging plant *Marchantia polymorpha* produces axes with non-photosynthetic scale-like lateral organs instead of leaves. Here we report that an ALOG (**A**rabidopsis **L**SH1 and **O**ryza **G**1) family protein in Marchantia, MpTAWAWA1 (MpTAW1), regulates meristem maintenance and lateral organ development. A mutation in Mp*TAW1*, preferentially expressed in lateral organs, induces lateral organs with mis-specified identity and increased cell number, and furthermore, causes defects in apical meristem maintenance. Remarkably, Mp*TAW1* expression rescued the elongated-spikelet phenotype of a rice mutant of Mp*TAW1* homologue. This suggests that ALOG genes are co-opted to specify lateral organ identities in both gametophyte and sporophyte shoots by repressing lateral organ growth. We propose that the recruitment of ALOG-mediated lateral organ modification was in part responsible for the convergent evolution of independently-evolved lateral organs among highly divergent plant lineages and contributed to the morphological diversification of land plants.

## Introduction

During 470 million years of evolution the body plans of land plants diversified independently among the gametophyte and sporophyte life stages of different plant groups. In extant bryophytes, basally-diverging land plants, the gametophyte is the dominant phase of the life cycle (1, 2). The gametophyte comprises an axis that develops from a meristem and forms structures in which gametes develop (antheridiophores and archaegoniophores). In contrast, the sporophyte is dominant in extant vascular plants. The sporophyte comprises an axial system (shoot or stem) that develops from a meristem and forms structures in which haploid spores develop. Therefore, despite their independent evolution, gametophytes and sporophytes develop axial systems that are produced by apical meristems (3-5).

Extant bryophytes and vascular plants develop lateral organs on gametophytes and sporophytes, respectively. Apical meristems maintain stem cell activity at their center and iteratively generate lateral organs at the meristem periphery. The spatio-temporal differences in cell division and expansion in lateral organs contribute to the morphological diversity of shoot structures in land plants (6-11).

The liverwort *Marchantia polymorpha* (*M. polymorpha*) is a bryophyte that forms an axis that undergoes indeterminate planar growth in the form of a flattened mat of tissue, called a thallus. The thallus exhibits strong dorsoventrality; gemma cups, gemmae and air chambers develop on the dorsal side while rhizoids and ventral scales are formed on the ventral side (Figure 1A) (12-17). Ventral scales cover bundles of rhizoids that run along the underside of the thallus and facilitate water and nutrient transport over the ventral surface of thallus (Figure 1B) (12). In the leafy liverworts, photosynthetic leaves arise next to a tetrahedral single stem cell (apical cell). By contrast, *M. polymorpha* does not develop photosynthetic leaves. Instead, the ventral scales alternately develop on the left and right sides of the wedge-shaped apical cell on the ventral surface in the apical notch near the growing tip of the thallus. The flattened form, single-cell thickness and bilateral symmetry resemble the leaves in leafy liverworts (Figure 1C) (12, 13). The ventral scales of *M. polymorpha* are hypothesized to be homologous to the photosynthetic leaves of the basally-diverging leafy liverworts (18, 19).

**Figure 1.**
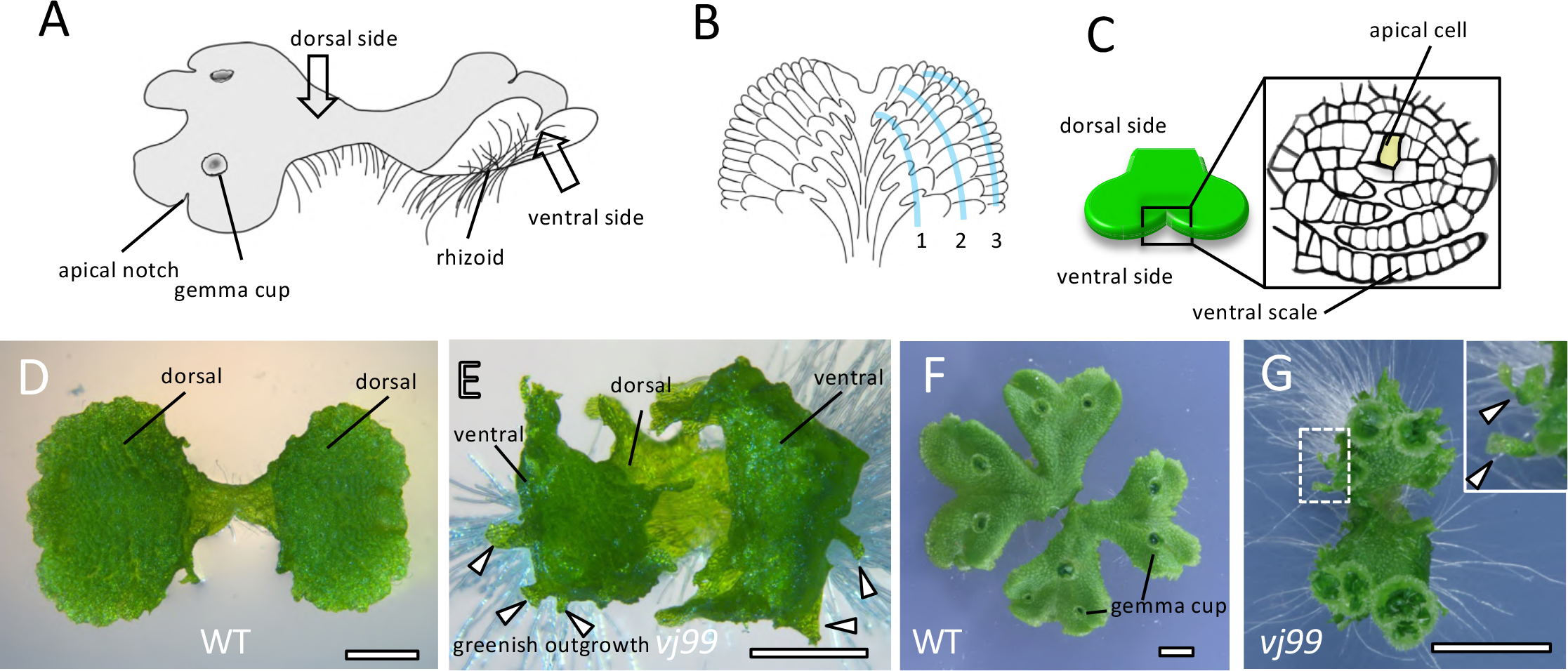
Illustration of *M. polymorpha* thallus and the isolation of Mp*taw1* mutants. (A-C) Diagrammatic representation of vegetative *M. polymorpha* thallus. Gross morphology (A), ventral side of thallus with ventral scales arranged in three rows on each side of the thallus (B), and vertical transverse section of a notch region (C). Rhizoids are not shown in (B) to clearly visualize ventral scales. The apical cell is shaded in (C). (D-G) Gross morphology of the wild-type (WT) Tak1 (D and F) and the Mp*taw1-1* mutant *vj99* (E and G) gemmalings. 10-d-old gemmalings (D and E) and 3-w-old (F and G) thalli are shown. The dotted square box in (G) that includes the abnormal green outgrowth is enlarged in the image in the top right corner. Arrowheads indicate abnormal green outgrowths. Note that unlike WT, Mp*taw1* mutants displayed upward bending of thalli and formed green outgrowths. Scale bars = 1 mm in (D and E), 0.5 cm in (F and G).

The fossil record indicates that the shoot of the earliest known land plants comprised branching stems without lateral, determinate organs (Kenrick and Crane, 1998). Subsequently, determinate lateral organs, which develop from the sides of apical meristems, evolved. The earliest example of such a lateral organ is the microphyll that developed on the stems of the sporophyte of *Baragwanathia longifolia*, a lycophyte, which first appears in the fossil record in the late Silurian (20, 21). No lateral organs are known from the gametophytes of early bryophytes from the Silurian or Devonian, but arose subsequently and are found in extant bryophytes. The acquisition and modification of different lateral organ types are likely to have been morphological adaptations to the terrestrial environment that increased photosynthetic efficiency, gas exchange and water transport (10).

Mechanisms controlling lateral organ development are well described in angiosperms such as rice and Arabidopsis. However, little is known about the mechanisms that regulate lateral organ development among bryophytes. Therefore, we carried out a forward genetic screen for mutants with defective lateral organ development in the liverwort *M. polymorpha* to define mechanisms that control lateral organ development in this species. Comparing the roles of the genes that control lateral organ development in liverworts and angiosperms will identify the mechanisms that were involved in the independent evolution of analogous lateral organs during land plant evolution.

## Results

### MpTAW1 specifies lateral organ identity during vegetative growth

We isolated two mutants, *vj99* and *vj86*, that produced abnormal green outgrowths from a population of 105,000 T-DNA transformed *M. polymorpha* (Figure 1D-1G, S1A and S1B). *vj99* and *vj86* thalli were hyponastic, bending upwards at the thallus margins unlike wild type (WT) (Figure 1D and 1E, S1C and S1D). A single T-DNA was inserted into the gene Mapoly0028s0118 in *vj99* and *vj86*, suggesting that defective function of Mapoly0028s0118 was responsible for the green outgrowths (Figure S1E). To test this hypothesis, we generated independent mutations in the Mapoly0028s0118 gene by homologous recombination. Mutants of Mapoly0028s0118 generated by targeted deletion developed similar phenotypes to those of the *vj99* and *vj86* mutants (Figure S1F and S1G). To verify that a defect in Mapoly0028s0118 was responsible for the green outgrowth we transformed mutant *vj99* with a genomic fragment that includes the full-length Mapoly0028s0118 gene. Transformation of the Mapoly0028s0118 genomic fragment into *vj99* mutants restored WT development, demonstrating that loss of Mapoly0028s0118 function causes the *vj99* phenotype (Figures S1H-K). Phylogenetic analysis indicated that Mapoly0028s0118 belongs to the ALOG (<u>A</u>rabidopsis <u>L</u>SH1 and <u>O</u>ryza <u>G</u>1) protein family (Figure S1L). The proteins in this family contain a DNA-binding domain with weak transcriptional activity (22-24). We named this gene Mp*TAWAWA1* (Mp*TAW1*) after the *TAWAWA1* (*TAW1*) gene in rice, an ALOG member that regulates meristem activity during reproductive growth (25). In addition to the abnormal green outgrowths, gemma cup spacing is abnormal in the Mp*taw1-1* (*vj99*) mutant; the distance between neighboring gemma cups is much shorter than the WT (Figure 1F and 1G).

To more precisely define the nature of the green outgrowths, we performed a phenotypic analysis of Mp*taw1-1* mutants. Outgrowths emerge from the ventral side of the thallus near the thallus margins and extend beyond the thallus margin in the Mp*taw1-1* mutant (Figure 2A-2D). These outgrowths resemble ventral scales in a number of ways. They develop in pairs near the apical notch (Figure 2E, 2F, S2A and S2B). They are in general a single cell layer thick, although the outgrowths located near the apical notch occasionally consist of several cell layers (Figure 2G-2I). Outgrowths located near the apical notch also tend to pile up on one another at the edge of the ventral surface (Figure 2G and 2I). Furthermore, while outgrowths develop, no ventral scales form on Mp*taw1-1* mutants (Figure 2C and 2D), suggesting that the outgrowths are modified ventral scales. Taken together, these data suggest that the outgrowths formed from the ventral thallus on Mp*taw1* mutants are related to ventral scales.

**Figure 2.**
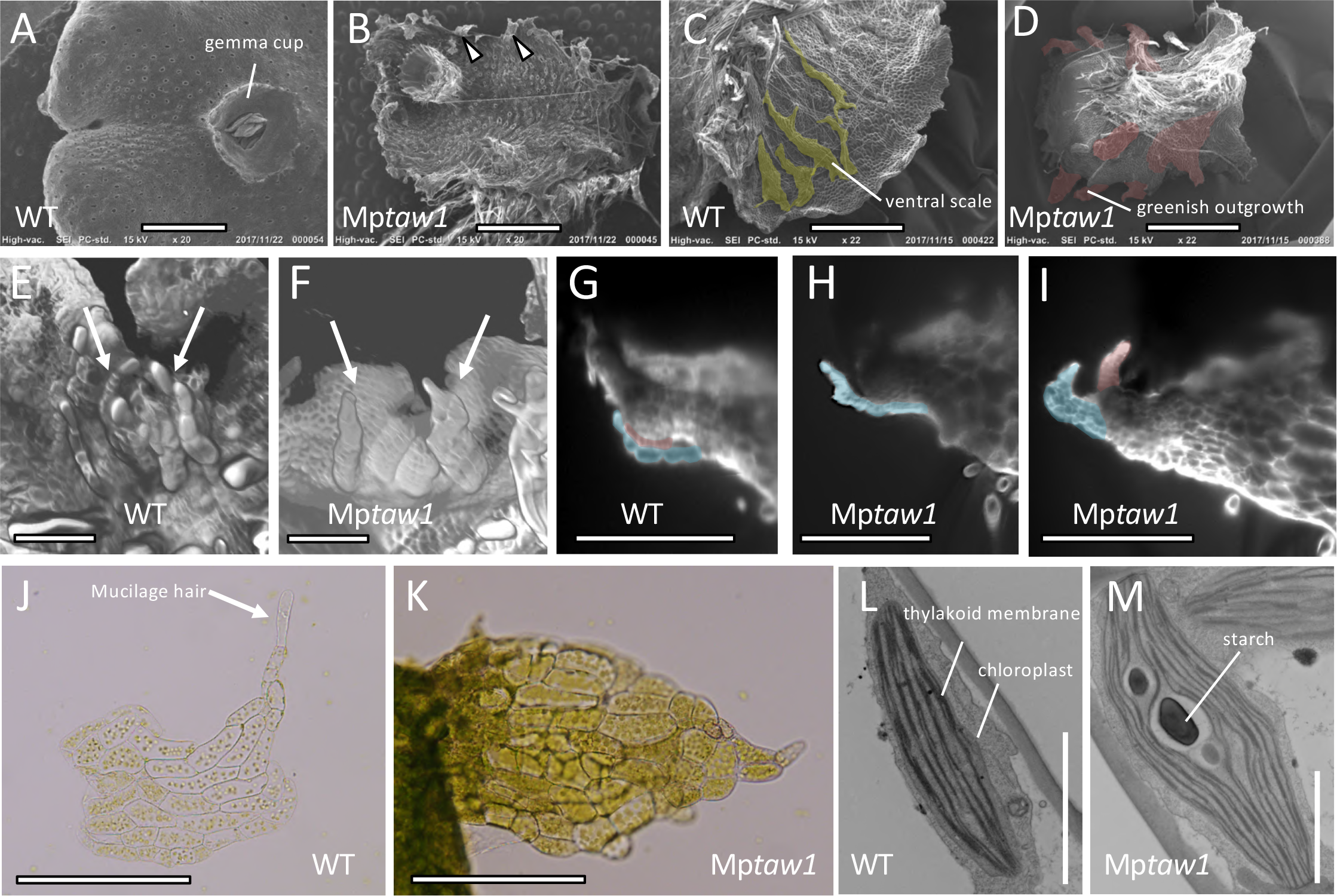
MpTAW1 functions are required for specification of lateral organs. (A-D) SEM images of WT (A and C) and the Mp*taw1* mutant (B and D) thalli. Dorsal (A and B) and ventral (C and D) sides of thalli are shown. Ventral scales in (C) and greenish outgrowth in (D) are highlighted in yellow and red, respectively. (E-I) LSFM (Light sheet fluorescence microscopy) images of WT (E and G) and Mp*taw1* mutants (F, H and I). Observation of apical notch regions from the ventral side (E and F) revealed the role of MpTAW1 in specifying lateral organs as ventral scales (F). Vertical longitudinal optical sections (G-I) around apical notches identified accelerated cell division of lateral organs in Mp*taw1* mutants (H and I). Lateral organs are indicated by arrows in (E and F) and highlighted in red or blue in (G-I). (J and K) Images of ventral scales in WT (J) and the corresponding tissues in Mp*taw1* mutants (K). Note that ventral scale cells are transformed into green tissues that lack mucilage hair cells in Mp*taw1* mutants. Mucilage hair in WT is indicated by an arrow in (J). (L and M) TEM images of ventral scale cells in WT (L) and the corresponding cells in Mp*taw1* mutants (M). Note that ventral scale cells are transformed into photosynthetic cells in Mp*taw1* mutants. Scale bars = 1.5 mm in (A-D), 150µm in (E and F), 300 µm in (G-I), 200 µm in (J and K) and 2 µm in (L and M).

Although similar to ventral scales, these outgrowths possess several substantially different characteristics. The abnormal outgrowths formed in Mp*taw1-1* mutants are greener than typical ventral scales (Figure 2J and 2K). There are more cells in outgrowths than in WT scales (Figures 2G-2I and S2C-S2F). Moreover, the mutant chloroplasts are larger than in WT and there are more thylakoid membranes in the mutants than in WT (Figure 2L and 2M). Starch, not seen in WT scales, accumulates in the green outgrowth in Mp*taw1-1* mutants, suggesting a higher photosynthetic activity in this tissue (Figure 2L and 2M). Rhizoids never differentiates in the green outgrowths of the Mp*taw1* mutants unlike wild-type ventral scales (Figures S2G and S2H). Taken together, these observations indicated that MpTAW1 plays crucial roles in specifying lateral organs as ventral scales, in which TAW1 inhibits cell division and chloroplast differentiation.

### MpTAW1 activity is required for the maintenance of meristem activity

The WT thallus comprises a bifurcating axis, and gemma cups develop along the midline of the dorsal surface. When the WT thallus axis bifurcates, a notch containing an apical cell forms on each of the two new axes (Figure 3A). The distance between each apical notch increases along with the forward growth of thalli. This process, the duplication of apical notches and the subsequent elongation of axes with separation of notches, is termed “axis separation”. Upon bifurcation, adjacent apical notches are initially pushed away by the growth of tongue-like tissues, named central lobes, and subsequently further separated concomitant with the axis elongation (Figure 3A) (26). Gemma cups initiate from dorsal merophytes, clones derived from the cell that are cut off from the dorsal surface of the apical cell (12), and they are regularly spaced along the dorsal midline of each axis. Gemma cups are more densely arranged along the dorsal surface of Mp*taw1* mutants than in WT (Figure 1G). This suggests defects in gemma cup differentiation or axis development, or both. To address whether Mp*TAW1* is involved in bifurcation or gemma cup differentiation, we analyzed the number of apices and gemma cups in Mp*taw1* mutants during cultivation. To count meristems we imaged expression of the *proYUC2A:GUS* construct that is preferentially expressed in notches (27). The number of apical notches expressing GUS was not significantly different between WT and Mp*taw1-1* mutants until day 7 of cultivation, although subsequently, less GUS-expressing apical notches were detected in the Mp*taw1-1* mutants compared to WT (Figure 3B-3D). This indicates that bifurcation occurs normally, at least in the early stages of development. In contrast, the density of apical notches in Mp*taw1-1* mutants was higher than WT at three weeks (Figure S3A and S3B). Importantly, there is no clear difference in the number of gemma cups between WT and Mp*taw1* mutants (Figure S3C). These data suggested that the onset of bifurcation, as well as gemma cup differentiation, are not affected, but that the separation process of each apical notch is compromised in Mp*taw1* mutants. The lower number of GUS-positive notches in Mp*taw1* mutants after 7 days of growth may be due to a secondary effect of slow thalli growth, or technical limitations of counting densely-clustered apices.

**Figure 3.**
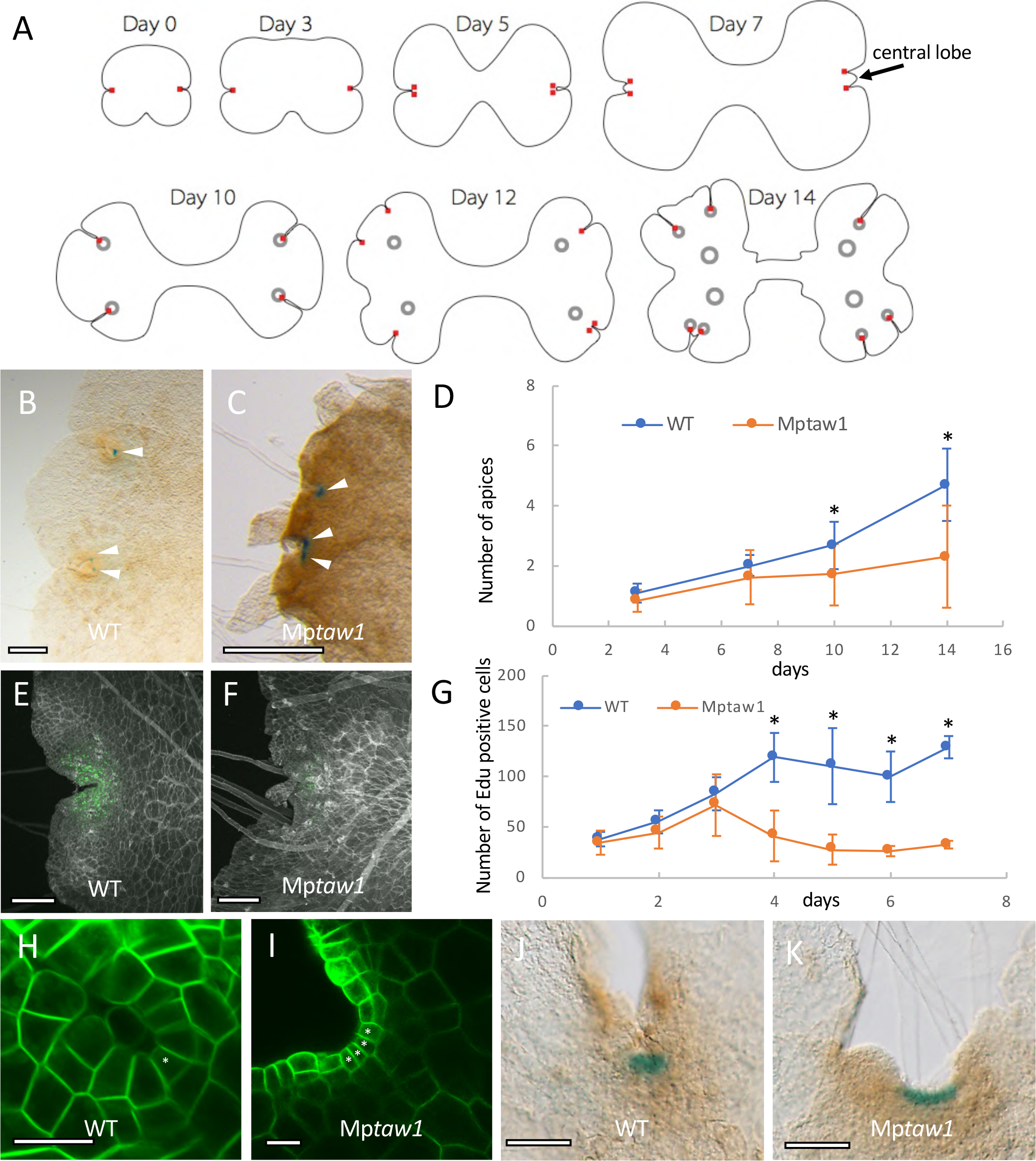
Loss-of-function mutant of Mp*TAW1* displays defects in meristem maintenance. (A) Diagrammatic representation of thallus shape transition in *M. polymorpha* thallus development. Note that distances between each apical notch as well as gemma cups are gradually increased along with the progression of bifurcation. Red squares and black circles indicate apical cells and gemma cups, respectively. (B and C) Apices stained by *proYUC2A:GUS* in 10-d-old gemmalings in WT (Tak1) (B) and Mp*taw1* mutants (C). Arrowheads indicate GUS staining at apical notches. (D) The number of apices stained by *proYUC2A:GUS* in Mp*taw1* mutants as compared to Tak1 throughout 14 d of gemmaling growth. The structure of a gemma is symmetrical, with a single notch on either sides. The number of apices originating from one side of each gemma (hereafter referred to as a “half gemmaling”) was counted. Each time point indicates the mean +/-SD. At least 14 gemmalings were analyzed at each time point. P values lower than 0.01 were indicated by asterisks (*). (E-G) Cell division activities in Mp*taw1* mutants decreased after 3 days’ incubation. EdU-positive signals of 4-d-old gemmalings in Tak1 (E) and Mp*taw1* mutants (F) are shown in green. (G) EdU uptake activities of Tak1 and Mp*taw1* mutants at the indicated time points. Edu-positive signals detected within half gemmalings were counted. Each time point indicates the mean +/-SD. At least 8 gemmalings were analyzed at each time point. P values lower than 0.01 are indicated by asterisks (*). (H-K) Defective apical meristem structures in Mp*taw1* mutants. A single apical cell, as seen in 3-d-old Tak1 gemmalings (H) was not observed in Mp*taw1* mutants (I). GUS staining of *proYUC2A:GUS* gemmalings in the apical notch region was broader in Mp*taw1* mutants (K) as compared to Tak1 (J). Plasma membranes (PMs) in (H) and (I) were labelled by *proEF:Lti6-GFP* constructs. Asterisks indicate triangular cells that are either apical cells or lateral merophytes. Scale bars = 500 µm in (B and C), 200 µm in (E and F), 20 µm in (H and I) and 100 µm in (J and K).

The separation of apical notches is dependent on the division and expansion of cells between notches (Figure 3A). We reasoned that defective cell division between the apical notches in Mp*taw1* mutant thalli leads to defects in apical notch separation. To test this possibility, we analyzed the cell division activity of Mp*taw1* mutant gemmalings during 7days’ cultivation by applying a 3 h pulse of 5-ethynyl-2’-deoxyuridine (EdU), a thymidine analog that is incorporated into cells during DNA replication (Figure 3E-3G)(28). The number of cells labelled by EdU was indistinguishable between the WT and Mp*taw1* mutants in 1-, 2- and 3-d-old gemmalings (Figures 3G). However, incorporation of EdU was lower in Mp*taw1* mutants than in WT between days 3 and 7 (Figures 3E-3G). This suggests that rates of DNA replication were lower in the mutant than in wild type, consistent with the hypothesis that cell division is reduced in the Mp*taw1* thallus compared to the wild type. We also compared the cellular organization of apical meristems of Mp*taw1* mutants and WT. The WT apical meristem comprises a single triangular apical cell and surrounding merophytes (cells derived from the apical cell), in which the lateral merophytes and the apical cell display identical shapes (Figure 3H)(12). In contrast, there are many triangular apical cells in Mp*taw1-1* mutants, in contrast to the single apical cell of wild type (Figure 3I). While the expression of *proYUC2A:GUS* is restricted to a small area of the WT apical notch, staining is more dispersed in Mp*taw1* mutants (Figures 3J and 3K). Occasionally (3 out of 20 gemmalings at 14 days’ cultivation) apical meristems are aborted in Mp*taw1* mutants (Figure S3D, dotted boxes), a phenomenon not observed in WT in our conditions. These data suggest that MpTAW1 is required for the maintenance of apical meristems. These data also support the hypothesis that Mp*taw1* mutants fail to separate apical notches due to defects in cell proliferation in the apical notches. Gemma cup differentiation as well as bifurcation initiate as in WT, but then subsequent defective meristem activity causes defective axis expansion, resulting in the development of a higher density of gemma cups and apical notches in the Mp*taw1* mutant thallus.

### MpTAW1 is expressed in lateral organs but not in apical cells

To define the spatial expression patterns of Mp*TAW1*, we established a line that expressed GUS under the control of 5’ and 3’ regulatory elements that were used in the complementation analysis of Mp*taw1* mutants (Figure S1E). In 4-d-old gemmalings, GUS staining was detected in notches and rhizoids (Figure 4A). Weak signal was observed elsewhere in growing thalli (Figures 4B and 4C). The developing ventral scales in the ventral region of the apical notch stained the strongest (Figures 4B-4F). Staining extended over the entire young ventral scale and the basal region of old ventral scales (Figures 4D and 4E). No signal was detected in the oldest ventral scales (Figures 4D and 4E). The expression of Mp*TAW1* in ventral scales is consistent with the phenotypic defects seen in these organs in the mutant, further strengthening our hypothesis that Mp*TAW1* is required for development of ventral scales. We also expressed functional *pro*Mp*TAW1:eGFP-*Mp*TAW1* constructs in Mp*taw1-1* mutants (Figures S1H, S1I and S1K) to analyse the distribution of MpTAW1 protein on a cellular level. eGFP-MpTAW1 protein was preferentially detected in the ventral parts of the apical notch regions (Figures 4G, 4H and S4; Supplementary Movie1). In particular, stronger signals were detected all over the young ventral scales as well as at the basal region of old ventral scales (Figures 4G, 4H and S4; Supplementary Movie1). These results further supported a crucial role for MpTAW1 in the specification of lateral organs as ventral scales. However, eGFP-MpTAW1 proteins were not detected in apical cells or lateral merophytes despite the defect in apical meristem morphology and maintenance in Mp*taw1-1* mutants (Figures 4G-4I). These findings suggest that MpTAW1 mediates the maintenance of apical meristems non-cell-autonomously, although we cannot exclude the possibility that MpTAW1 proteins below the level of detection in the apical meristems maintain meristem activity (Figure 4I).

**Figure 4.**
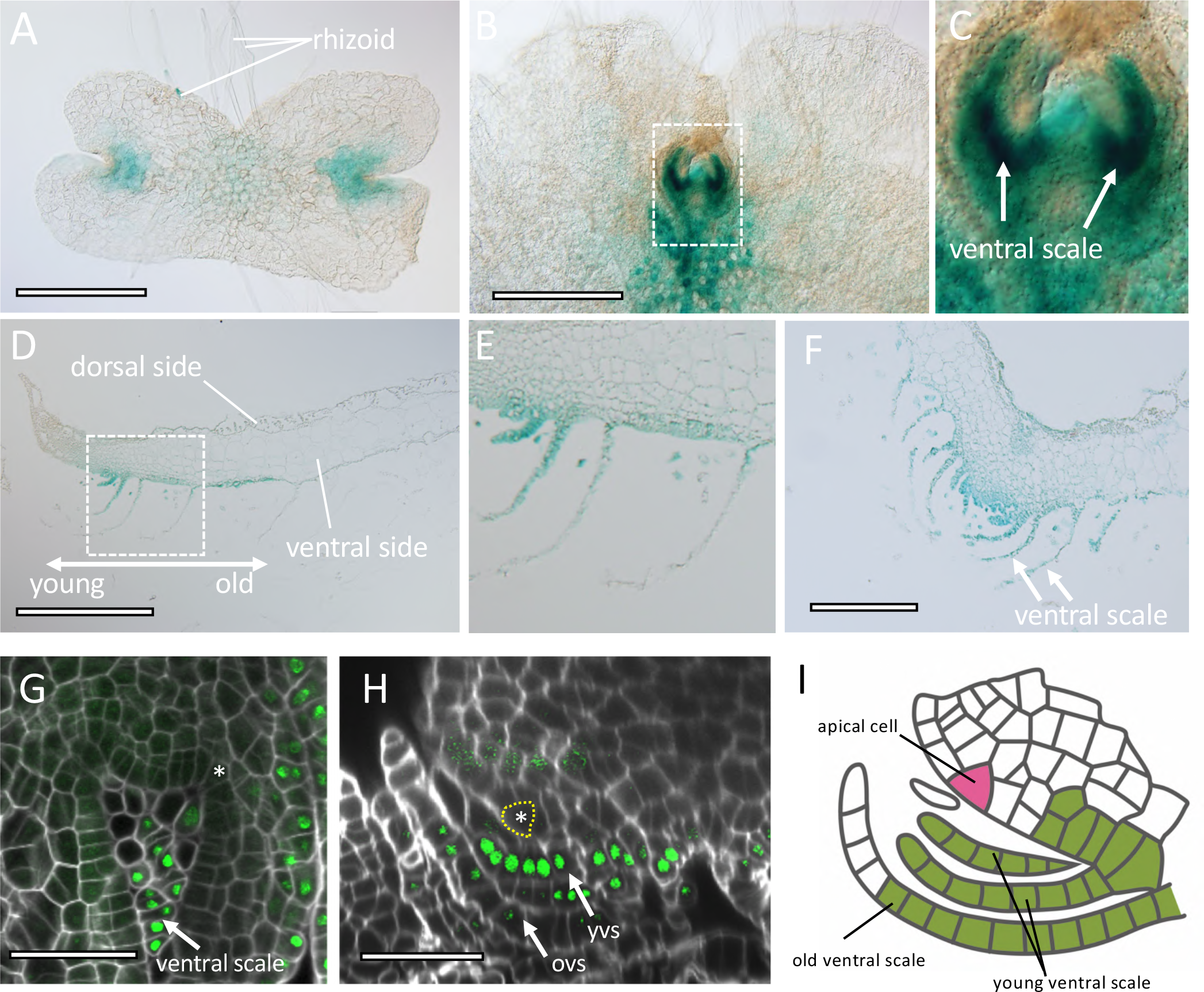
MpTAW1 regulates lateral organ development cell-autonomously and meristem maintenance non-cell-autonomously. (A-C) GUS activity in ventral thalli of 4-d-old (A) and 10-d-old (B and C) *pro*Mp*TAW1:GUS*-expressing gemmalings. (C) is a close-up image of the dotted square depicted in (B). (D-F) Cross-section of GUS-stained 10-d-old *pro*Mp*TAW1:GUS* gemmalings. (D and E) are vertical longitudinal sections and (F) is a vertical transverse section. (E) is a close-up image of the dotted square depicted in (D). “young” and “old” describe in (D) indicates the relative age of ventral scales. (G and H) Functional eGFP-MpTAW1 proteins were not detected in apical meristems but were detected in ventral scales and the cells beneath the basal part of ventral scales. Confocal images of a horizontal optical section (G) and a vertical longitudinal optical section (H) of Mp*taw1* mutant gemmalings that express *proMpTAW1:eGFP-*Mp*TAW1* constructs are shown. Cell walls were stained by calcofluor. Apical cells are indicated by asterisks and/or dotted yellow lines. “yvs” and “ovs” indicates young ventral scales and old ventral scales, respectively. (I) Schematic of MpTAW1 protein localization around apical notches. MpTAW1 proteins are detected at ventral scales and the cells that are located around the basal part of ventral scales. Note that MpTAW1 protein is not present in apical cells. Green indicates cells that contain MpTAW1 protein. The apical cell is in pink. Scale bars = 500 µm in (A, B and D), 200 µm in (F) and 50 µm in (G and H).

### MpTAW1 specifies lateral organ identity during reproductive growth

*M. polymorpha* produces an umbrella-like gametangiophore (antheridiophore or archegoniophore) that bears antheridia or archegonia during reproductive growth (12). The gametangiophore is a vertically growing thallus branch (12) and we reasoned that gametangiophore development might be defective in Mp*taw1-1* mutants. The antheridial receptacles of male Mp*taw1-1* plants are smaller than WT and, unlike in the WT, antheridia are frequently exposed (Figures S5A-S5C). Moreover, the scales on the antheridial receptacles of Mp*taw1-1* plants are larger than WT (Figures S5D-S5H). Mp*TAW1* expression was detected in the ventral scales of the antheridiophore, as well as jacket cells, mucilage cells, and throughout the antheridia in plants harboring the *proMpTAW1::GUS* transgene (Figures S5I and S5J). These observations demonstrate that MpTAW1 regulates ventral scale development by restricting cell division in both the vegetative and reproductive phases.

The archegonial receptacle of female *M. polymorpha* is highly lobed, with finger-like structures called digitate rays (Figure 5A). The archegonial receptacle lacks the rows of typical ventral scales that develop in antheridiophores. Instead, a pair of specialized scale-like structures called involucres, which are larger than ventral scales, develop between each digitate ray and enclose the archegonia cluster (Figures 5C and 5D)(12). In female Mp*taw1-2* mutants, large leaf-like structures developed as in antheridiophores (Figure 5E). Moreover, more than two involucre-like structures differentiated between each digitate ray (Figures 5C-5F). Importantly, these involucre-like structures resemble ventral scales in their arrangement in several rows (Figures 1B, 1C, 5G and 5H). This suggests that loss of MpTAW1 function results in the homeotic transformation of involucres into more scale-like structures. GUS staining was also detected in immature involucres but not in mature involucres in *proMpTAW1::GUS* archegoniophores (Figures 5I and S5K), accompanied by staining of all parts of the archegonia including eggs, collars and venters (Figures 5I, S5K and S5L). These data suggest that ventral scales are transformed into involucres in a MpTAW1-dependent manner upon the transition from vegetative to reproductive growth, in which MpTAW1 inhibits the growth of two rows of ventral scales, resulting in the formation of a single pair of involucres.

**Figure 5.**
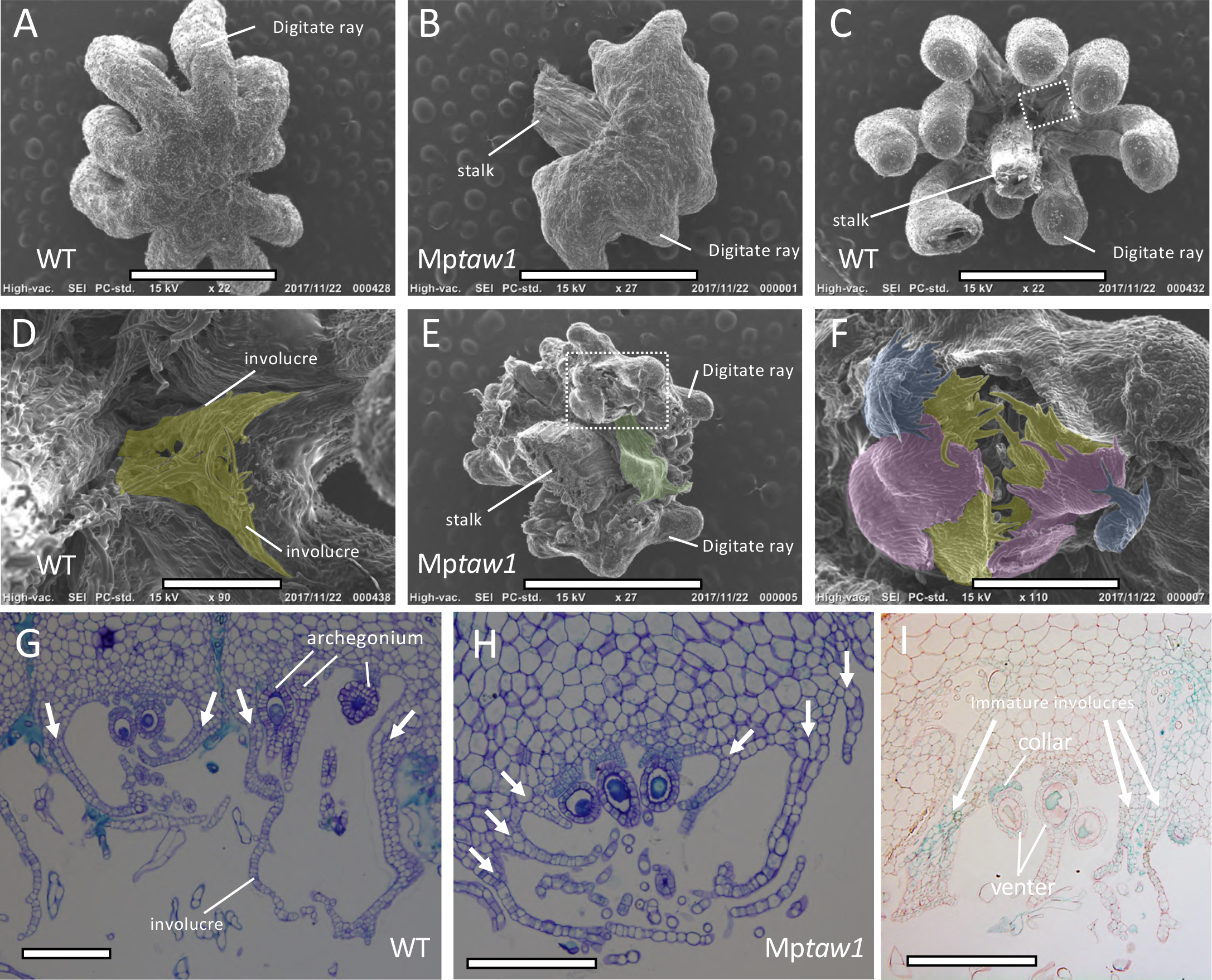
MpTAW1 is necessary for the specification of lateral organ identity during reproductive growth. (A and B) SEM images of the dorsal side of archegoniophores in WT Tak2 (A) and in Mp*taw1* mutants (B). Note that archegoniophores in Mp*taw1* mutants display shorter, malformed finger-like structures in place of digitate rays. (C-F) SEM images of the ventral side of archegoniophores in WT (C and D) and in Mp*taw1* mutants (E and F). (D) and (F) are magnified images of (C) and (E), respectively. Instead of a single pair of involucres as developed by the WT (C and D), three pairs of membranous structures developed in Mp*taw1* mutants (E and F). Enlarged leaf-like structures (highlighted in green) also developed in Mp*taw1* mutants (E). Involucres in WT and the three pairs of membranous structures in Mp*taw1* mutants are highlighted in yellow, red or blue. (G and H) Cross-sections of archegonial receptacles at regions between the digitate rays in WT (G) and in Mp*taw1* mutants (H). Note the three pairs of ventral scale-like membranous structures in Mp*taw1* mutants. Arrows indicate involucres or ventral scale-like structures. (I) Cross-sections of GUS-stained archegoniophores that express *pro*Mp*TAW1:GUS.* GUS activity was detected in immature involucres. Scale bars = 2 mm in (A, B, C and E), 400 µm in (D and F) and 200 µm in (G, H and I).

### Molecular function of ALOG proteins are conserved between Marchantia and rice

Mutation of *G1*, a member of the *ALOG* gene family in rice, results in the enlargement of sterile lemmas; a small leafy lateral organ in the rice spikelet (the basic unit of a grass flower), which is interpreted as a homeotic transformation of a sterile lemma into a lemma (22). Similarly, Mp*taw1* mutants display defects in lateral organ specification, that we interpret as transformation of involucres into ventral scales. To determine whether the *M. polymorpha* protein could rescue the homeotic transformation of the rice mutant, we expressed Mp*TAW1* in rice *g1* mutants. Expression of Mp*TAW1* restored the WT short sterile lemma phenotype (Figure 6A-6C). This suggests that the molecular functions of the ALOG proteins have been conserved since the time that *M. polymorpha* and rice last shared a common ancestor, which likely lacked lateral organs. It further suggests that ALOG family proteins were independently co-opted to specify sporophytic function in the lineage giving rise to rice and gametophytic functions in the lineage giving rise to liverworts, when each originated the evolutionary novelty of lateral organs.

**Figure 6.**
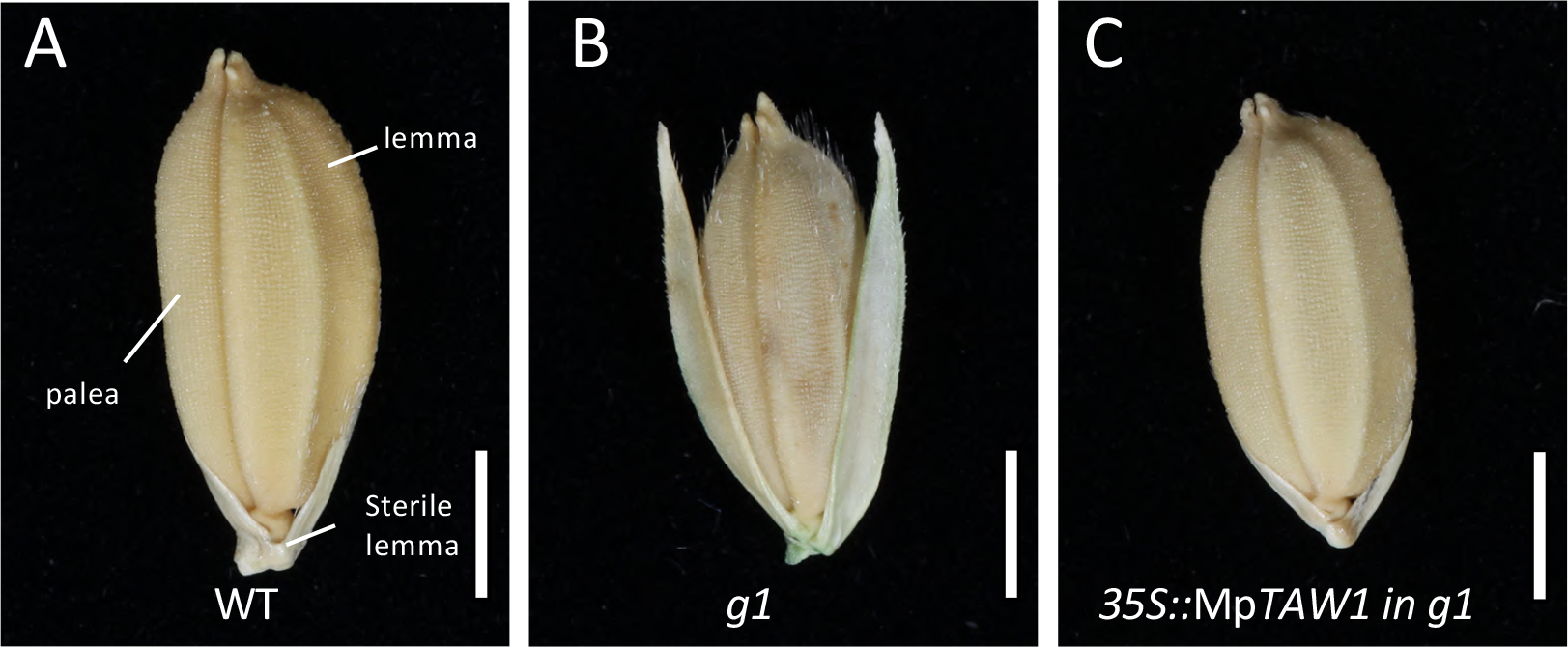
Evolutionarily-conserved ALOG family proteins in Marchantia and in rice are co-opted to specify analogous lateral organs. (A-C) Complementation of rice *alog* mutants with Mp*TAW1*. Phenotypes of a WT (A), a *g1* mutant (B) and a *g1* mutant expressing the *35Spro::*Mp*TAW1* construct (C) are shown. Scale bars = 2 mm in (A, B and C).

## Discussion

### Land plant ALOG proteins regulate lateral organ development and meristem activity

Here we report the discovery that MpTAW1 controls both lateral organ development and apical meristem activity in *M. polymorpha*. MpTAW1 represses the growth of different lateral organs, including ventral scales and involucres, and Mp*TAW1* expression was detected early in the development of these lateral structures. These data indicate that the gene is required for lateral organ development. Furthermore, MpTAW1 activity is required for apical meristem maintenance. However, Mp*TAW1* is not expressed in the apical meristems or surrounding cells. We propose that MpTAW1 cell-autonomously regulates lateral organ development but non-cell-autonomously regulates apical meristem maintenance (Figure 4I).

The role of TAW1 proteins in meristem maintenance is conserved between monocots and dictos. OsTAW1 and SlTMF (the tomato TAW1 homolog) proteins repress maturation of meristems during reproductive growth (23, 25, 29). While the angiosperm genes control meristem development, neither SlTMF, AtLSH3 (the Arabidopsis TAW1 homolog) nor OsTAW1 proteins are expressed in apical meristems. Instead they are expressed at lateral organ boundaries (24, 29, 30). Taken together, these data from a diversity of land plants suggest that while the ALOG genes act cell-autonomously during the development of lateral organs, they act non-cell-autonomously to control meristem development. It remains unclear how this might operate, but there is evidence from angiosperms that lateral organ development is required for meristem maintenance (8, 31, 32).

Taken together with our discovery that MpTAW1 is required non-cell-autonomously for meristem maintenance in *M. polymorpha*, this means that the evolutionary-conserved ALOG family proteins control apical meristems in divergent plant lineages, in which the apical meristems are found in different phase of the life cycle. We propose that this mechanisms for controlling shoot meristematic activity was already present in the last common ancestor of Marchantia and rice, the earliest land plants.

### Conserved ALOG proteins negatively regulating lateral organ growth

We discovered that MpTAW1 specifies lateral organ identity by negatively regulating the lateral organ outgrowth; involucres are transformed into ventral scale-like structures during reproductive growth in Mp*taw-1* mutants (Figures 5G and 5H). We interpret the transformation of involucres into scales as a homeotic transformation, in the same way that loss-of-function mutations in a homologous gene in rice result in homeotic transformations in the spikelet. The rice homolog, G1 also represses the development of lateral organs to specify the sterile lemmas. Loss-of-function mutations in *OsG1* result in the transformation of small sterile lemmas into large lemmas (Figures 6A and 6B). This transformation of one member of a meristic series into another member is designated a homeotic transformation (33). Similarly, *Sltmf* mutants develop a similar homeotic transformation where sepals develop leaf characteristics (29). The conserved function of rice, tomato and *M. polymorpha* TAW1 homologs suggests that the role in lateral organ repression, as evidenced by homeotic transformation in loss-of-function mutations, is ancient. This functional conservation among divergent taxa of land plants suggests two alternative hypotheses regarding the evolution of lateral organs. According to the first hypothesis, the ALOG gene controlled the development of lateral organs in the last common ancestor of the liverworts and the seed plants. These structures subsequently diverged morphologically during the course of land plant evolution. An alternative hypothesis is that the ALOG-dependent growth-repression mechanism existed in the last land plant common ancestor which lacked lateral organs. The ALOG mechanism was subsequently recruited independently during the evolution of lateral organs in different lineages leading to the liverworts and seed plants. Both hypotheses are consistent with our discovery of the role for ALOG genes in land plants. However, the currently best-supported hypotheses based on the fossil record is that the last common ancestor of liverworts and seed plants lacked lateral organs and instead developed naked shoot axes (34, 35). Lateral organs subsequently evolved independently in different land plant lineages. If the last common ancestor did not develop lateral organs and since ALOG function regulates lateral organ development in both liverworts and seed plants, we suggest that ALOG function was recruited independently during the evolution of lateral organs in different lineages of land plants. The recruitment of ALOG function during the independent evolution of lateral organs provides a molecular mechanism for the convergent evolution of lateral organs.

### ALOG proteins may mediate diversification of lateral organs during plant evolution

It has been suggested that the repressive activity of the *OsG1* gene on growth led to the evolution of the rice spikelet (22). Loss-of-function *Osg1* mutations revert the sterile lemma into a larger leafy structure which has been interpreted as similar to a hypothetical ancestral structure (Figures 6A and 6B) (22). According to this model, the formation of a pair of lower lemmas subtending the floret (rice flower surrounded by two bracts; the external lemma and internal palea) was the ancestral state. Then, during the evolution of rice, *OsG*1 activity was co-opted to repress the development of the lower lemma, which is now much reduced in size in modern rice compared to the ancestral state, resulting in the formation of the sterile lemma. It is formally possible that *MpTAW1* may also have played a similar role in the evolution of lateral organs in liverworts. Several liverwort taxa with thalloid form are suggested to have evolved independently from ancestral leafy liverworts, where leaves are hypothesized to be transformed into non-photosynthetic ventral scales with reduced growth during this evolutionary transition (18, 19). We found that Mp*TAW1* is involved in the specification of lateral organ identities by inhibiting cell division and chloroplast differentiation, and that the loss of its function leads to the formation of chlorophyll-containing photosynthetic tissues (Figures 2J and 2K). These green appendages are in fact similar to the green-colored photosynthetic scales formed in the Treubiaceae family of liverworts, whose semi-thalloid form has been interpreted as an evolutionary transition state between the leafy and thalloid form (36-38). Therefore, MpTAW1 function may also be associated with the evolution of thallose body form by repressing leaf growth in ancestral leafy liverworts in the same way that gene Os*G1* suppresses lower lemma development during rice spikelet evolution. It is possible that morphological modification of lateral organs is controlled by the spatial and temporal differences in expression levels of ALOG family genes and this would provide the mechanism for the establishment of morphological diversification in lateral organs that develop on shoots during land plant evolution.

### Conclusion

We demonstrate that MpTAW1 plays a function in integrating meristem activity and lateral organ differentiation in *M. polymorpha*. MpTAW1 acts by repressing lateral organ growth and is required for meristem maintenance. Since ALOG proteins from angiosperms also repress lateral organ growth and are required for meristem maintenance, and these functions were rescued by MpTAW1, we conclude that molecular functions of ALOG family proteins are conserved between among these taxa and acted in their last common ancestor. We hypothesize that ALOG genes were co-opted to execute the morphological modification of analogous lateral organs during land plant evolution and contributed to diversification of lateral organs in shoot systems during the course of land plant evolution.

## Material and Methods

### Plant materials and growth conditions

*Marchantia polymorpha* Takaragaike-1 (Tak1, male) and Takaragaike-2 (Tak2, female) gemmalings were grown for 3–60 days at 22 °C under continuous light on petri plates containing 1/2 Gamborg’s Basal Salt Mixture (B-5) growth medium at pH 5.5 with 1.2 % agar (Nacalai tesque, Japan). Transition to reproductive growth was induced through far-red light supplementation (39).

### Generation of mutant plants

Knockout mutants of Mp*taw1-1 (vj99)* and Mp*taw1-2 (vj86)* were isolated by a mutant screen of spores from a cross between Tak1 and Tak2 transformed with the T-DNA vector pCambia1300 as previously described (14, 40). Knockout mutants of Mp*taw1-3* and Mp*taw1-4* were generated by gene-targeted homologous recombination (41). Genetic nomenclature is outlined in (42).

### Phylogenetic analysis

Phylogenetic analysis was performed as described by Bowman et al. (5). Protein sequences were collected using the Marchantia genome portal site MarpolBase (http://marchantia.info). Multiple sequence alignments were performed using the MUSCLE program (43) contained in the Geneious software (https://www.geneious.com). Gaps were removed by using Strip Alignment Columns in the Geneious package and phylogenetic analyses were performed using PhyML (http://www.atgc-montpellier.fr/phyml/).

### Plasmid construction

For constructing the *pro*Mp*TAW1:*Mp*TAW1* plasmid that complements the Mp*taw1* mutants, an Mp*TAW1* genomic fragment with a 10 kb upstream region and a 3 kb downstream region was amplified by PCR using Prime STAR GXL polymerase (TaKaRa, Japan) and subcloned into pENTR/D-TOPO (Thermo Fisher Scientific, USA), which was subsequently integrated into pMpGWB101 by a Gateway LR reaction (44). pENTR/D-TOPO that included the *pro*Mp*TAW1:*Mp*TAW1* complement fragment was modified to establish *pro*Mp*TAW1:eGFP-*Mp*TAW1* and *pro*Mp*TAW1:*Mp*TAW1-GUS* plasmids. A PCR-amplified *eGFP* coding sequence was inserted in frame with the 5’ end of the MpTAW1 coding sequence by the In-Fusion cloning reaction (TaKaRa, Japan) to generate the *pro*Mp*TAW1:eGFP-*Mp*TAW1* plasmid. The coding sequence of MpTAW1 in the *pro*Mp*TAW1*:Mp*TAW1* complement fragment was replaced with a PCR-amplified GUS coding sequence by the In-Fusion cloning reaction to generate *pro*Mp*TAW1:*Mp*TAW1-GUS* plasmids. pENTR/D-TOPO vectors that included *pro*Mp*TAW1:eGFP-*Mp*TAW1* or *pro*Mp*TAW1:*Mp*TAW1-GUS* fragments were subsequently integrated into pMpGWB101 by the Gateway LR reaction.

### Histochemical GUS staining

GUS staining was performed as described by Naramoto et al. (45), except that 50 mM sodium phosphate buffer was used. Samples were cleared with 70 % ethanol and subsequently mounted using a clearing solution (chloral hydrate:glycerol:water, 8:1:2) for direct microscopic observation or dehydrated through a graded ethanol series and embedded in paraffin or Technovit 7100 resin for microtome sectioning.

### Plant embedding and sectioning

Plant material was fixed in FAA (45 % ethanol:5 % formaldehyde:5 % acetic acid in water) for embedding in paraffin and Technovit 7100. For paraffin embedding, fixed plant material was dehydrated in a series of ethanol (25–50 %), t-butyl alcohol (10–75 %) and chloroform (20 %) solutions, then embedded in Paraplast (McCormick, USA). For Technovit 7100 embedding, fixed sample was dehydrated through a graded ethanol series and embedded in Technovit 7100 resin, according to the manufacturer’s instructions (Heraeus Kulzer, Germany). Embedded samples were sectioned on a rotary microtome into a series of vertical transverse and longitudinal sections (thickness of 8 μm for paraffin and 4 μm for technovit sectioning). The obtained sections were further treated with neutral red dyes as a counterstain for GUS-stained samples or toluidine blue for the other samples. Multi-Mount 480 solution (MATSUNAMI, Japan) or Entellan new (MERCK, USA) were used as mounting agents to preserve the samples on the slides.

### ClearSee treatment and staining of cell walls

Plants were fixed with 4 % paraformaldehyde (PFA) in 1× PBS for 1 h at room temperature under vacuum. Samples were subsequently washed twice with PBS and transferred to ClearSee solution (10 % xylitol, 15 % sodium deoxycholate and 25 % Urea in water) (46). ClearSee treatment was prolonged until samples became transparent. Cell walls were stained for 1 h with 0.1 % (v/v) Calcofluor or with 0.1 % (w/v) Direct Red23, dissolved in ClearSee. Stained samples were washed for at least 30 min with ClearSee solution before observation.

### EdU uptake experiments

Gemmalings were incubated in 1/2 B5 medium containing 10uM EdU (Click-iT Edu Alexa Fluor 488 imaging kit; Thermo Fisher, USA) for 3h. Samples were fixed with 4 % PFA in 1 × PBS for 1 h under vacuum and then washed three times in PBS. Coupling of EdU to the Alexa Fluor substrate was performed according to the manufacturer’s instructions. Before observations, samples were cleared with ClearSee solution and cell walls were subsequently stained by Direct Red 23.

### Microscopy

Anatomical features were observed with a light microscope (Olympus BX51) equipped with an Olympus DP71, a light sheet microscope (Zeiss Z.1) or a confocal laser scanning microscope (Olympus FV1000 or Zeiss LSM880). For light microscope observations, a PLAPON 2x objective, a UPlanFl 10× objective or a UPlanFl 20× objective were used. Light sheet microscope observations were conducted using Lightsheet Z.1 detection optics 5× or Clr Plan-Neofluar 20×. For confocal laser scanning microscopy, cell walls stained by Calcofluor or by Direct Red23 were excited at 405 nm or 543 nm, respectively, whereas GFP and Alexa 488-labelled EdU were excited at 488 nm. Samples were mounted using ClearSee solution and observed with silicon oil objectives. The 3D reconstruction was done by using Imaris software (BITPLANE, http://www.bitplane.com/). High-resolution images showing ultrastructural details were obtained using a Scanning Electron Microscope (JEOL JCM-6000Plus NeoScope) and Hitachi SU820.

## Acknowledgements

We thank Kei Saito, Eriko Kida and Kanane Sato for assistance with transformation and microtome sectioning. We also thank Mayumi Wakazaki for preparing samples for electron microscopy. This work is supported by Grants-in-Aid from the Ministry of Education, Culture, Sports and Technology, Japan (KAKENHI grant numbers 17K17595 for S.N., 18H04836 for R.N., 17H06472 for K.I. and 17H06465 for J.K.). V.A.S.J. was funded by a Newton Abraham Studentship from the University of Oxford. LD was funded by an ERC advanced Grant EVO-500 (250284).

## Author contributions

S.N. carried out the majority of experiments. V.J. and L.D. isolated *vj99* and *vj86* mutants. S.N. and N.T. conducted SEM and GUS analyses. M.S. and K.T. conducted TEM analyses. S.N., V.J., M.S., S.I., K.N., K.I., R.N., T.K. L.D, and J.K. analyzed data. S.N., V.J., M.S., L.D., and J.K. designed the project. S.N., V.J., M.S., L.D., and J.K. wrote the manuscript.

## Supplementary Figures

**Figure S1. Responsible gene of *vj99* mutant encodes an ALOG family protein (related to Figure 1).**

(A and B) Phenotypes of *vj99* and *vj86* mutants.

(C and D) LSFM image of gross morphology of WT (C) and *vj99* mutant (D) gemmalings.

(E) Overview of the functional Mp*TAW1* construct and the T-DNA insertion mutants isolated by forward genetic screening. The regions that harbor 10152 bp upstream and 1689 downstream of coding sequences were used to express Mp*TAW1*.

(F and G) Phenotypic series of Mp*taw1* knockout mutants.

(H-K) Complementation of Mp*taw1* mutants with a functional Mp*TAW1* construct. The phenotype of Mp*taw1-1* (I) is complemented by introducing the genomic Mp*TAW1* fragment (J) as well as the *eGFP*-fused Mp*TAW1* genomic fragment (K).

(L) Phylogenetic tree of ALOG family proteins in Arabidopsis, rice and Marchantia. Green, red and blue symbols indicate ALOG proteins in Arabidopsis, rice and Marchantia, respectively.

Scale bars = 1cm in (A, B, F, G, H, I, J and K), 500 µm in (C and D).

**Figure S2. The mutation in Mp*TAW1* transforms ventral scales into chlorophyll-containing photosynthetic tissues with an increased number of cells. (related to Figure 2).**

(A and B) LSFM image of gemmalings in WT (A) and in Mp*taw1* mutants (B) observed from the ventral side. Dotted boxes in (A) and (B) that include apical notch regions are shown as close-up images in Figure 2 (E) and (F), respectively. Ventral scales in WT or their corresponding tissues in Mp*taw1* mutants are indicated by arrows.

(C-F) Vertical transverse optical sections of apical notch regions in 4-d-old gemmalings obtained by LSFM. Optical sections in WT (C and D) and in Mp*taw1* mutants (E and F) are shown. Note that the number of cells that comprise mucilage and ventral scales increased and thus these tissues became larger in Mp*taw1* mutants.

(G and H) Images of ventral scales in WT (G) and the corresponding tissues in Mp*taw1* mutants (H). Note that ventral scale cells are transformed into green tissues that lack rhizoids and mucilage hair cells in Mp*taw1* mutants. Rhizoids and mucilage hair in WT are indicated by an arrowhead and an arrow, respectively.

Scale bars = 150 µm in (A and B), 100 µm in (C and E) and 200 µm in (G and H).

**Figure S3. MpTAW1 plays crucial roles in maintenance of meristem activities (related to Figure 3).**

(A and B) Apices stained by *proYUC2A:GUS* in 3-w-old-gemmalings in WT (A) and Mp*taw1* mutants (B). Arrowheads indicate GUS staining at apical notches. (C) Number of gemma cups in WT Tak1 and in Mp*taw1* mutants. Each bar indicates the mean +/-SD. At least 7 gemmalings were analyzed at each time point.

(D) SEM image of Mp*taw1* mutant gemmalings. Note the frequent termination of meristem activity which accompanies thalli regeneration in Mp*taw1* mutants. The meristems presumed to be aborted are indicated by dotted boxes.

Scale bars = 1.5 mm in (A and B) and 1 mm in (D).

**Figure S4. Functional eGFP-MpTAW1 proteins were detected in ventral scales and the cells beneath the basal part of the ventral scales (related to Figure 4).**

Confocal laser scanning microscopy (CLSM) image of Mp*taw1* mutant gemmalings that express *pro*Mp*TAW1:eGFP-*Mp*TAW1* constructs. Vertical transverse sections were obtained after the 3D reconstruction of a series of CLSM images. Cell walls were stained by calcofluor.

Scale bar = 50 µm.

**Figure S5. MpTAW1 plays crucial roles in lateral organ differentiation in reproductive growth (related to Figure 5).**

(A-F) SEM image of antheridiophores in WT Tak1 (A and D) and Mp*taw1* mutants (B, C, E and F). The dorsal side (A-C) and ventral side (D-F) of antheridiophores are shown.

(C) and (F) are close-up images of (B) and (E), respectively. Some ventral scales are highlighted by colors. Note the exaggerated growth of ventral scales as compared to the size of thalli (F).

(G and H) Vertical sections of antheridia in WT (G) and Mp*taw1* mutants (H). Note the extra cell division of mis-specified ventral scales in Mp*taw1* mutants. Ventral scales (G) and mis-specified ventral scales (H) are indicated by arrows.

(I and J) Cross-sections of GUS-stained antheridiophores (I) and antheridia (J) that express *pro*Mp*TAW1:GUS* constructs. Note that GUS activities were detected in the ventral scales in WT (I).

(K and L) Cross-sections of GUS-stained archegoniophores that express *pro*Mp*TAW1:GUS.* Regions that include mature involucres (K) and whole image of archegoniophores (L) are shown. Note that in contrast to immature archegoniophores, GUS activity was not detected in mature involucres.

Scale bars = 2 mm in (A,B and E), 1mm in (F), 400 µm in (C), 200 µm in (G,H,I,K and L).

**Supplemental Movie 1. 3D reconstruction of the apical notch region by using CLSM data that display eGFP-MpTAW1 and cell walls.**

Cell walls were stained by calcofluor.

